# The Integrin Binding Peptide, ATN-161, as a Novel Therapy for SARS-CoV-2 Infection

**DOI:** 10.1101/2020.06.15.153387

**Authors:** Brandon Beddingfield, Naoki Iwanaga, Prem Chapagain, Wenshu Zheng, Chad J. Roy, Tony Y. Hu, Jay Kolls, Gregory Bix

## Abstract

Many efforts to design and screen therapeutics for severe acute respiratory syndrome coronavirus (SARS-CoV-2) have focused on inhibiting viral cell entry by disrupting ACE2 binding with the SARS-CoV-2 spike protein. This work focuses on inhibiting SARS-CoV-2 entry through a hypothesized α5β1 integrin-based mechanism, and indicates that inhibiting the spike protein interaction with α5β1 integrin (+/− ACE2), and the interaction between α5β1 integrin and ACE2 using a molecule ATN-161 represents a promising approach to treat COVID-19.

---

As of June 26, 2020, there have been 484,249 deaths out of a total 9,473,214 confirmed COVID-19 cases, for an estimated fatality rate of 5.5% (https://www.who.int/emergencies/diseases/novel-coronavirus-2019). This viral outbreak began in China in late 2019 (2), with a likely origin in bats, with selection resulting in efficient human-to-human transmission occurring before or after transfer to the human host (3). This follows the same epizoontic transmission events seen in other severe viral infections, including SARS-CoV (4) and Ebola (5), and was predicted prior to this outbreak (6). Interaction between the SARS-CoV-2 spike protein and the angiotensin-converting enzyme II (ACE2) receptor has been implicated in SARS-CoV-2 entry and replication (7). Many therapeutic efforts spurred by the current pandemic have focused on disrupting an aspect of the viral replication process (8, 9), including host entry (10), often focusing on inhibition of ACE2/spike protein binding (11).

Integrin binding has also been implicated in the SARS-COV-2 cell entry mechanism, as the spike protein contains an integrin binding motif (RGD) (12–16). Integrins are extracellular matrix receptors expressed throughout the body, including in the respiratory tract (e.g. epithelial cells (17)) and vasculature (e.g. endothelial cells (18)), and the β1 family of integrins are closely associated (in proximity and functional regulation) with ACE2 (19, 20). A non-RGD peptide derived from the extracellular matrix component fibronectin, referred to herein as ATN-161, can bind to and inhibit the activity of certain integrins, including α5β1 (21, 22), and has been previously used to study viral replication (23). ATN-161 binds outside the RGD-binding pocket, thus acting as a non-competitive inhibitor of integrin binding, especially for α5β1 (24). Likewise, ACE2 binds to α5β1 in an RGD-independent fashion, although it possesses an RGD motif in a region inaccessible for protein-protein interaction (19, 20).

Molecular docking of ATN-161 with ACE2 or ACE2-spike RBD complex revealed three potential biding sites as shown in Fig 1A. One of these is at the interface between the ACE2 and the spike RBD. This may affect the binding of RBD with the ACE2. ATN-161 is also found to bind the integrin alpha5beta1 ectodomain complex near the RGD motif binding site located at the interface between the α5 and β1 chain (25), potentially affecting the binding of α5β1 with proteins containing the RGD motif. Although ACE2 contains the RGD sequence, it is inaccessible for binding under physiological conditions. Therefore, it is believed that another sequence KGD (residues 353, 354,355), which closely resembles the sequence RGD, may bind α5β1 via the RGD-binding site (26). Figure 1B shows the ACE2-α5β1 complex obtained from protein-protein docking using Zdock with the ACE2 residues around KGD and the α5β1 residues around the RGD-binding site selected as preferred binding partners. This docking results in a complex with the desired orientations of the integrin chains (27) and ACE2 relative to the plasma membrane (Fig 1A). As shown in Fig. 1B, the binding of the α5β1 to ACE2 at this site masks the binding site for the spike RDB, potentially inhibiting the SARS-CoV-2 entry (26). The binding of ATN-161 in the interface may disrupt the α5β1-ACE2 complex.

**Figure 1.**
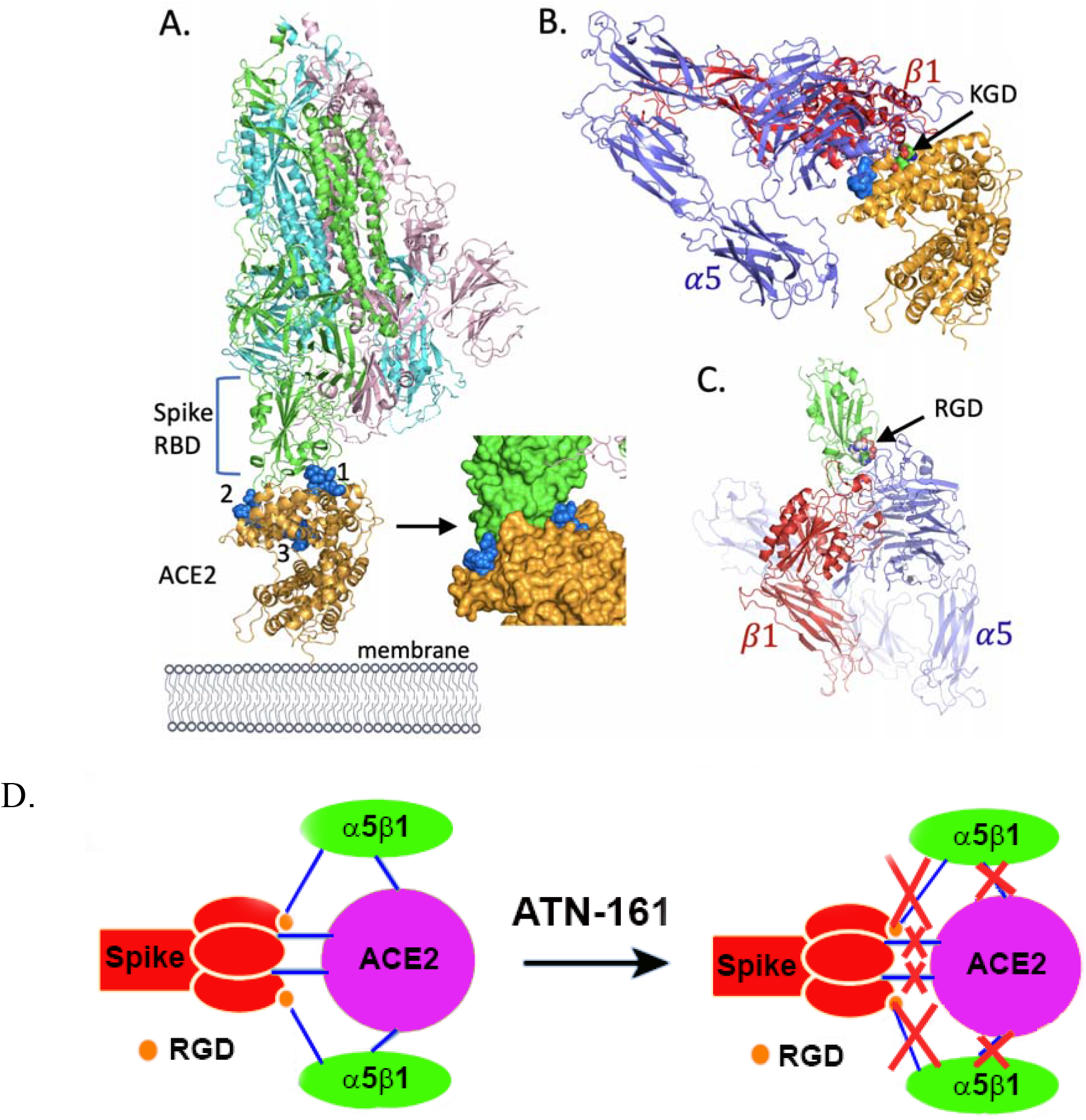
Molecular Model of ATN-161 Interactions with α5β1. (A) SARS-CoV-2 spike protein trimer bound to ACE2 via the sprung out spike protein receptor binding domain (RBD). Molecular docking of ATN-161 shows three potential binding sites (van der Waals representation in blue). (B) ACE2-alpha5beta1complex, with the KGD sequence is highlighted. The location of ATN-161 in site 2 is highlighted (blue surface representation) but was not included for protein-protein docking. (C) The Spike RBD-alpha5beta1 complex, with RGD sequence of the spike RBD highlighted. All conformations have the same orientation relative to the membrane in A. (D) Proposed mechanism of ATN-161 inhibition of SARS-CoV-2 infection, where addition of ATN-161 is proposed to inhibit SARS-CoV-2 spike protein binding to host α5β1 integrin, ACE2, as well as α5β1-ACE2 binding.

Separately, we performed docking of α5β1 to the spike protein RBD, which contains the RGD sequence that is accessible for binding. This results in a complex of the spike RBD and α5β1 as shown in Fig. 1C. For this binding to occur, the RGD-binding interface of integrin needs to be oriented differently than binding with ACE2, consistent with the active conformation of integrin (7). ATN-161 binding near the RGD motif binding site of integrin may inhibit the α5β1-spike RBD complex formation. We hypothesize that SARS-CoV-2 entry is facilitated by binding to ACE2-associated α5β1 integrin via its spike protein, and that ATN-161 treatment will inhibit infection by blocking this binding event and by disrupting the initial ACE2 and α5β1 interaction (Figure 1D).

In this study, we explored the binding of the SARS-CoV-2 spike protein with ACE2 and α5β1, utilizing ELISA-based methods. To determine the spike protein’s ability to bind α5β1, plates were coated with α5β1 and incubated with a mixture of ATN-161 and a trimeric version of the spike protein. The SARS-CoV-2 spike protein binds to α5β1 with an affinity that is roughly equivalent to α5β1’s native ligand, fibronectin (28), and inhibits binding with a U-shaped (Donate et al., 2008) dose-dependent manner, with maximum effect at 100nM (Figure 2A). To our knowledge, this is the first report of SARS-CoV-2 spike protein interaction with integrins, and specifically α5β1. We performed similar assays to investigate ACE2 binding to α5β1, using a mixture of ATN-161 and human ACE2 protein (hACE2). Clear inhibition of ACE2/α5β1 binding by ATN-161 was apparent and dose-dependent (Figure 2B). Furthermore, application of ATN-161 reduced binding of trimeric spike protein to hACE2, either alone or in combination with α5β1, the latter of which trended to support greater spike binding than to hACE2 alone (Figure 2C). Application of ATN-161 also reduced binding of monomeric spike to hACE2 (Figure S1).

**Figure 2.**
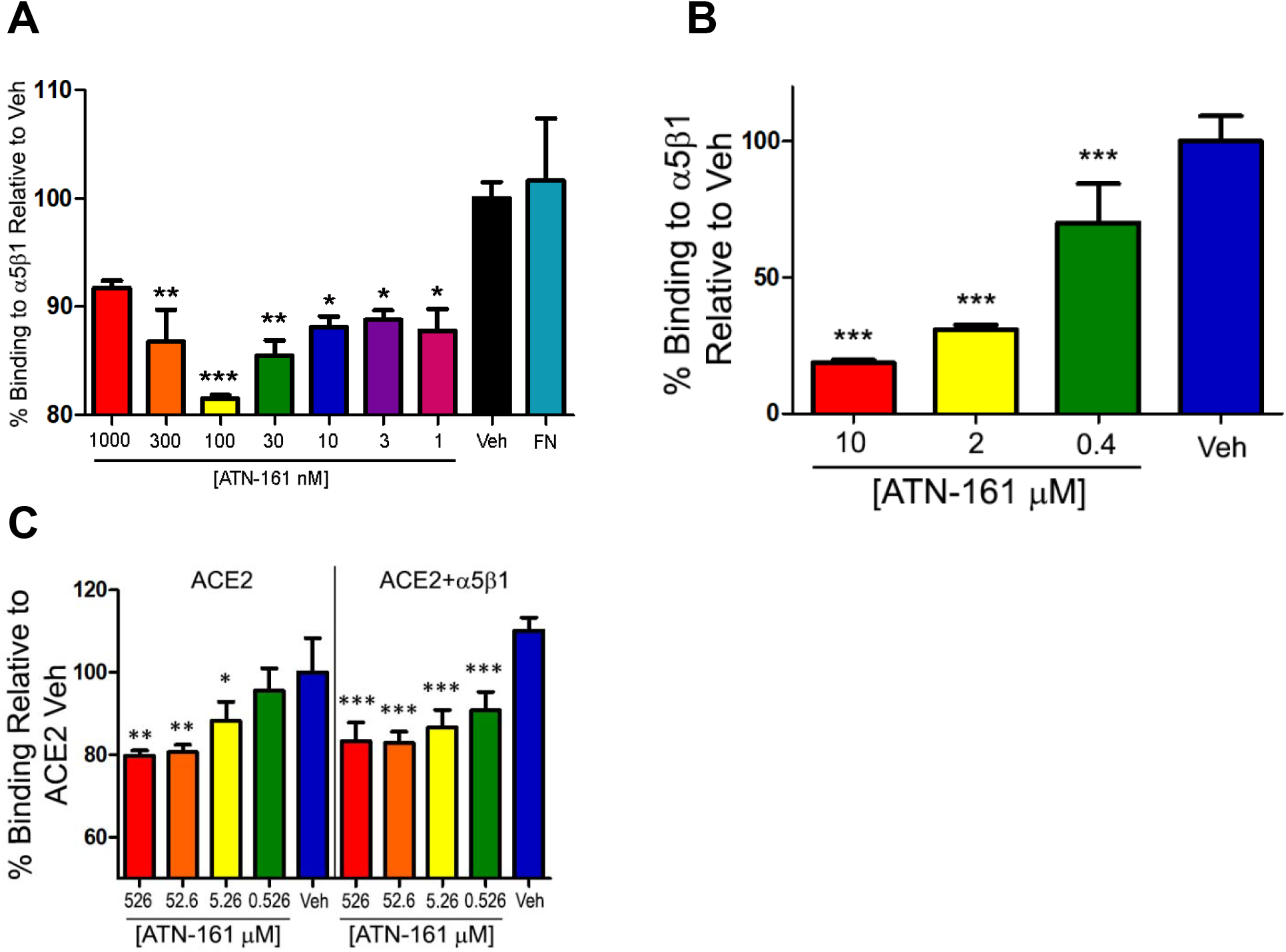
ATN-161 effects on SARS-CoV-2 spike, ACE2 and α5β1 binding. ELISA data indicates that ATN-161 alters (A), binding of α5β1 to spike protein-coated plates, when these plates are incubated with α5β1 and concentrations of ATN-161 and (B) when α5β1-coated plates are incubated with human ACE2 and concentrations of ATN-161, and (C) spike binding to ACE2 or ACE2 + α5β1 protein-coated plates. Data was normalized to no-ATN vehicle control (ACE2 Veh for D, stats as compared to respective Veh). Data represent mean ± SD, n=3, * P<0.05, ** P<0.01, *** P<0.001

The *in vitro* assessment of ATN-161 and therapeutic potential was performed using a once-passaged Vero (E6) African green monkey (*Chlorocebus atheiops*) kidney cell line utilizing competent SARS-CoV-2. ATN-161 was effective at reducing viral loads after infection (Figure 3A), with an estimated IC_50_ of 3.16 μM. The EC_50_ value for ATN-161 approximates the value for remdesivir (8). Importantly, Vero (E6) have been previously shown to express α5β1 integrin (29).

**Figure 3.**
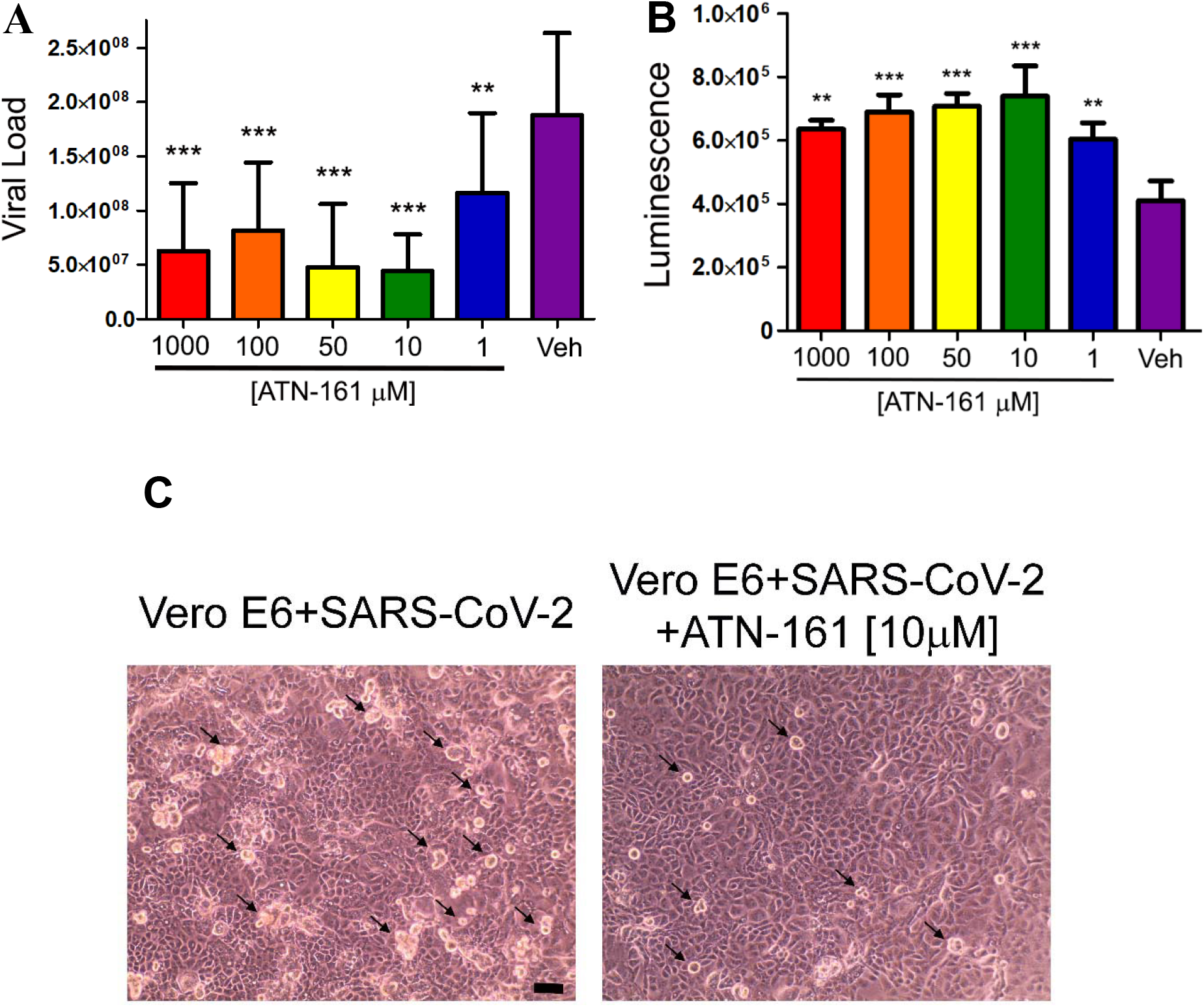
ATN-161 effects on SARS-CoV-2 infection *in vitro*. (A) Viral Load of SARS-CoV-2 with and without ATN-161 treatment. VeroE6 cells were incubated with indicated ATN-161 concentrations for one hour prior to 48 hours infection with SARS-CoV-2 at a MOI of 0.1. (B) Cell viability via luminescence-based CellTiterGlo 24 hours post-infection. (C) Representative phase contrast microscope images of VeroE6 cells 24 hours post-infection with and without 10 μM ATN-161 treatment. Black arrows indicate some of the visible viral cytopathic effect (rounded, phase bright cells). Scale bar is 10 μM. Data represent mean ± SD, n=3, * P<0.05, ** P<0.01, *** P<0.001

Measuring cellular viability and underlying cytotoxicity is another metric for antiviral therapeutic potential that we explored with ATN-161 (30). After 24 hours infection at a MOI of 0.01, cells were lysed with CellTiterGlo and luminescence values were taken to measure ATP production in each treatment. Pretreatment with ATN-161 increased ATP production in infected cells, indicating increased viability, and was consistent with viral PCR data at concentrations as low as 1μM ATN-161 (Figure 3B). Addition of 10 μM ATN-161 resulted in a decreased cytopathic effect (i.e. fewer apparent rounded, bright cells) when cells were visualized by phase contrast microscopy (Figure 3C).

In summary, we show that SARS-CoV-2 spike protein binds to both α5β1 and α5β1/hACE2, and that this binding can be effectively inhibited by ATN-161, which also disrupts SARS-CoV-2 infection *in vitro*. Prophylactic treatment of ATN-161 increased cell viability in the presence of SARS-CoV-2 and decreased cytopathic effects associated with viral infection. Taken together, and in light of ATN-161’s previously demonstrated *in vivo* therapeutic efficacy against a closely related beta-coronavirus (porcine hemagglutinating encephalomyelitis virus (23)) and its successful use in human cancer clinical trials (31), these results support the performance of *in vivo* studies to assess the potential efficacy of ATN-161 as an experimental therapeutic agent for COVID-19.

## Methods

### Cells and Virus

VeroE6 cells (ATCC# CRL-1586) were cultured in complete DMEM containing 10% fetal bovine serum (FBS). SARS-CoV-2 stock from viral seed (SARS-CoV-2; 2019-nCoV/USA-WA1/2020 (BEI# NR-52281) was obtained by infecting nearly confluent monolayers of VeroE6 cells for one hour with a minimal amount of liquid in serum free DMEM. Once adsorption was complete, complete DMEM containing 2% FBS was added to the cells and the virus was allowed to propagate at 37□ in 5% CO_2_. Upon the presence of CPE in the majority of the monolayer, the virus was harvested by clearing the supernatant at 1,000 xg for 15 minutes, aliquoting and freezing at −80□. Sequencing confirmed consensus sequence was unchanged from the original isolate.

### ELISA Analysis of ATN-161 Inhibition of SARS-CoV-2 Spike Protein Binding to ACE2 and Integrin

Enzyme-Linked Immunosorbent Assay (ELISA) was utilized to determine the ability of ATN-161 to disrupt binding events essential to entry of SARS-CoV-2 into a host cell. For determination of inhibition of ACE2/ α5β1 integrin binding by ATN-161, α5β1 was coated on 96-well plates at 1 μg/mL for 2 hours at room temperature and blocked overnight with 2.5% BSA. Addition of 0.5 μg/mL of hACE2-Fc (Sino Biological, Cat# 10108-H02H) in differing concentrations of ATN-161 followed, incubating for 1 hour at 37□. Incubation with an HRP labeled goat anti-human Fc secondary antibody at 1:5000 for 30 minutes at 37□ was followed by detection by TMB substrate.

In order to assess disruption of binding of α5β1 to SARS-CoV-2 Spike protein, 96-well plates were coated as before, but incubation with ATN-161 was performed in conjunction with 1μg/mL spike (produced under HHSN272201400008C and obtained through BEI Resources, NIAID, NIH: Spike Glycoprotein Receptor Binding Domain (RBD) from SARS-Related Coronavirus 2, Wuhan-Hu-1, Recombinant from HEK293 Cells, NR-52306) in the presence of 1mM MnCl_2_, followed by detection with an anti-spike antibody. The rest of the procedure was consistent with the previous.

### *In vitro* assessment of ATN-161 Inhibition of SARS-CoV-2 Infection

In order to determine the ability of ATN-161 to reduce the infection capability of SARS-CoV-2 *in vitro*, a cell-based assay was utilized. VeroE6 cells were plated at a density of 1.25 × 10^4^ cells/well in a 96-well plate and incubated overnight at 37□ in 5% CO_2_. The next day, cells were treated with dilutions of ATN-161 in complete DMEM with 2% FBS for one hour at 37□ in 5% CO_2_, followed by viral infection at an MOI of 0.1. After 48 hours, virus and cells were lysed via Trizol LS and RNA was extracted using a Zymo Direct-zol 96 RNA Kit (#R2056) according to manufacturer’s instructions. Experiments were performed under Biosafety Level 3 conditions in accordance with institutional guidelines.

### RT-qPCR

Viral load was quantified using a Reverse Transcriptase qPCR targeting the SARS-CoV-2 nucleocapsid gene. RNA isolated from cell cultures was plated in duplicate and analyzed in an Applied Biosystems 7300 using TaqPath supermix with the following program: i)50□ for 15 min., ii) 95□ for 2 min. and iii) 45 cycles of 95□ for 3s and 55□ for 30s. The primers and probes were as follows: 2019-nCoV_N1 Forward :5’-GAC CCC AAA ATC AGC GAA AT-3’, 2019-nCoV_N1 Reverse: 5’-TCT GGT TAC TGC CAG TTG AAT CTG-3’, and 2019-nCoV_N1 Probe: 5’-FAM-ACC CCG CAT TAC GTT TGG TGG ACC-BHQ1-3’. Standard curves were generated for each run using a plasmid containing SARS-CoV-2 nucleocapsid gene (Integrated DNA Technologies, USA).

### Cell Imaging

The day before infection, Nunc LabTek II chamber slides (Thermo, USA) were seeded with 2.5 × 10^4^ cells per chamber. On the day of infection, chambers were treated with dilutions of ATN-161 in complete DMEM with 2% FBS for one hour prior to infecting with SARS-CoV-2 at an MOI of 0.01. Slides were placed in a 37□ 5% CO_2_ incubator for 24 hours prior to imaging via phase contrast using an EVOS XL inverted microscope (Thermo, USA).

### Cell Viability Assay

Ability of ATN-161 to increase cell viability was performed with CellTiterGlo (Promega, USA). Cell supernatant was removed 24 hours post infection and cells were lysed via pre-mixed CellTiterGlo reagent. Cells were incubated for 15 minutes and allowed to shake briefly before ATP was quantified via luminescence readout on the GloMax Explorer multimode plate reader (Promega).

### Molecular Modeling

The structure of the ACE2-Spike protein receptor binding domain complex (7) was obtained from the protein data bank (PDB ID 6m0j). To get the orientation of the SARS-CoV-2 spike protein trimer relative to ACE2, the receptor binding domain (RBD) was aligned with the sprung out RBD of the prefusion conformation of the spike protein trimer (PDB ID 6vsb) (32). Similarly, the integrin α5β1 ectodomain structure (25) was obtained from the protein data bank (PDB ID 3vi3), with the calf1 and calf2 domains of α5 added from the PDB ID 6naj (33). ATN-161 (Ac-PHSCN-NH2) was prepared for docking with Autodock vina (34). ATN-161 was docked α5β1 complex, ACE2, and ACE2-spike RBD complex. ZDock 3.0.2 (35) server was used for protein-protein docking to generate the α5β1 complexed with ACE2 as well as with the spike RBD. The structures were rendered using PyMol 2.3.0 (36).

### Statistics

Differences between groups was determined via the one-way ANOVA using Dunnett’s multiple comparisons test. Experiments are represented as weighted mean and standard deviation of a total of three replicates. For IC_50_ estimation, the data points directly bounding the IC_50_ value were used and calculation was made in GraphPad Prism. Viral load studies were performed 3 separate times with each condition done in triplicate in each experiment. All ELISA studies were performed two times with each condition done in triplicate. The cell viability assay was performed a single time, each condition in triplicate.

## ACKNOWLEDGEMENT

We thank R Garry, Tulane University School of Medicine, for use of PCR reagents, N Maness at Tulane National Primate Research Center for viral stock, A Mazar, Monopar Therapeutics, for helpful discussions on the preparation and handling of ATN-161 for in vitro studies, I Rutkai and A Narayanappa for collection of technical information regarding antibodies used in ELISA studies, and S Yadav for advice on 3D modeling. We would also like to thank K Andersen at Scripps Research Institute for sequencing of viral stock. G.B. is supported by Tulane University startup funds. JKK was supported by the following NIH grant R35HL139930 for this work. This research was supported in part by grant OD0011104 to CJR from the National Center for Research Resources and the Office of Research Infrastructure Programs (ORIP), NIH. TYH was supported by Department of Defense grant W8IXWH1910926 and NIH grants R21EB026347, R01AI122932, R01AI113725, R01HD090927 and R21AI126361.

## AUTHOR CONTRIBUTIONS

G.B. conceived the study. B.B. conducted all live SARS-CoV-2 studies. N.I. and W.Z. conducted all ELISA’s. B.B., N.I. and W.Z. collected data and performed computational analysis. P.C. performed 3D modeling analysis. B.B., C.R., T.H., J.K. and G.B. interpreted data and wrote the manuscript with input from all of the authors.

---

**Fig. S1.**
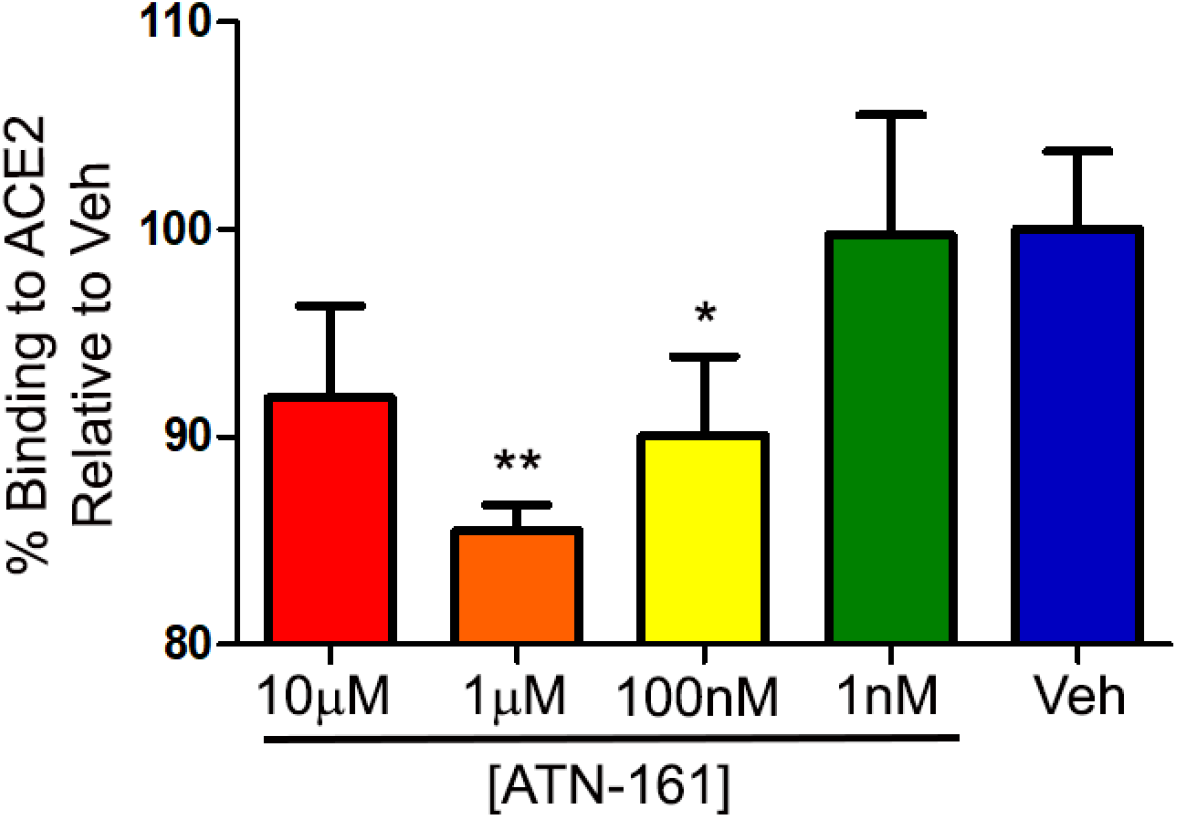
Inhibition of SARS-CoV-2 spike protein binding to human ACE2 by ATN-161. Plates were pre-coated with monomeric spike protein and incubated with a mixture of hACE2 and various ATN-161 concentrations, followed by detection of bound hACE2 via HRP-conjugated anti-ACE2 antibody. Data was normalized to a no-ATN vehicle control. Data represent mean ± SD, n=3, * P<0.05, ** P<0.01.

